# Stem cell differentiation is a stochastic process with memory

**DOI:** 10.1101/101048

**Authors:** Patrick S. Stumpf, Rosanna C. G. Smith, Michael Lenz, Andreas Schuppert, Franz-Josef Müller, Ann Babtie, Thalia E. Chan, Michael P. H. Stumpf, Colin P. Please, Sam D. Howison, Fumio Arai, Ben D. MacArthur

## Abstract

Pluripotent stem cells are able to self-renew indefinitely in culture and differentiate into all somatic cell types *in vivo*. While much is known about the molecular basis of pluripotency, the molecular mechanisms of lineage commitment are complex and only partially understood. Here, using a combination of single cell profiling and mathematical modeling, we examine the differentiation dynamics of individual mouse embryonic stem cells (ESCs) as they progress from the ground state of pluripotency along the neuronal lineage. In accordance with previous reports we find that cells do not transit directly from the pluripotent state to the neuronal state, but rather first stochastically permeate an intermediate primed pluripotent state, similar to that found in the maturing epiblast in development. However, analysis of rate at which individual cells enter and exit this intermediate metastable state using a hidden Markov model reveals that the observed ESC and epiblast-like ‘macrostates’ conceal a chain of unobserved cellular ‘microstates’, which individual cells transit through stochastically in sequence. These hidden microstates ensure that individual cells spend well-defined periods of time in each functional macrostate and encode a simple form of epigenetic ‘memory’ that allows individual cells to record their position on the differentiation trajectory. To examine the generality of this model we also consider the differentiation of mouse hematopoietic stem cells along the myeloid lineage and observe remarkably similar dynamics, suggesting a general underlying process. Based upon these results we suggest a statistical mechanics view of cellular identities that distinguishes between functionally-distinct macrostates and the many functionally-similar molecular microstates associated with each macrostate. Taken together these results indicate that differentiation is a discrete stochastic process amenable to analysis using the tools of statistical mechanics.

## Introduction

Two distinct pluripotent states are found in the pre-gastrulation mouse embryo (Nichols & Smith 2009): a naïve pluripotent state that emerges from the inner cell mass of the blastocyst between E3.5 and E4.5 (Evans & Kaufman 1981, Martin 1981) and a primed pluripotent state that emerges after implantation of the blastocyst into the uterus wall at E5.5 (Tesar et al. 2007, Brons et al. 2007). During this naïve-to-primed pluripotency transition, cells undergo dramatic changes to their signaling requirements, transcriptional regulatory control mechanisms and global epigenetic status (Nichols & Smith 2009). These molecular changes are accompanied by morphological changes of the pluripotent tissue *in vivo* (Tam & Loebel 2007). Following this transition, cells become increasingly susceptible to the spatially coded differentiation cues that determine the foundation of the principal germ layers in the body. A variety of molecular mechanisms regulate this susceptibility in order to prevent premature lineage commitment and enable the correct formation of the egg-cylinder, including the regionalisation of the extra-embryonic endoderm and hence the foundation for the differential signaling gradients across the embryo during gastrulation (Tam & Loebel 2007). At this stage, the timely release of pluripotency maintenance mechanisms is just as important as the gain of lineage-specific characteristics (Betschinger et al. 2013, Nichols & Smith 2009, Turner et al. 2014), and appropriate differentiation is regulated by the balance of these two processes. However, despite recent interest in this problem (Moris et al. 2016, Semrau et al. 2016, Hormoz et al. 2016) the dynamics of exit from the pluripotent state at the individual cell level are only partially understood. In particular, while it is known that stochastic fluctuations in key transcription factors have an important role in the early stages of differentiation (Chambers et al. 2007, Toyooka et al. 2008, Hayashi et al. 2008, MacArthur & Lemischka 2013, Abranches et al. 2014) it is not yet clear if cellular responses to these fluctuations are also stochastic, or if this inherent molecular stochasticity is buffered such that differentiation progresses in a deterministic way through a continuum of intermediary cell states (MacArthur et al. 2012, Moris et al. 2016, Semrau et al. 2016, Hormoz et al. 2016, Trapnell et al. 2014, Bendall et al. 2014). To address these questions we sought to profile a well-defined transition in detail in order to elucidate the dynamics of individual cells in transition from the pluripotent state to a specified committed lineage.

## Results

### Differentiation *in vitro* recapitulates developmental dynamics *in vivo*

Starting from the pluripotent ground state in LIF + 2i conditions (Ying et al. 2008), the closest in *vitro* equivalent to the naïve pluripotent state of the pre-implantation epiblast (Boroviak et al. 2014, 2015, Kolodziejczyk, Kim, Tsang, Ilicic, Henriksson,Natarajan, Tuck, Gao, Bühler, Liu, Marioni & Teichmann 2015), we directed differentiation of mouse embryonic stem cells (ESCs) in mono-layer culture towards the neuroectoderm using a well-established protocol (Ying et al. 2003) that utilizes the neuroinductive property of retinoic acid (Bain et al. 1996, Lu et al. 2009, Stavridis et al. 2010, Rhinn & Dollé 2012), a metabolite of vitamin A contained in N2B27 medium (**Fig. 1a**). This transition was chosen since it has previously been shown to induce robust and reliable differentiation (Ying et al. 2003, Lu et al. 2009, Abranches et al. 2009, Kelava & Lancaster 2016), and therefore serves as a good model system to examine the kinetics of the exit from pluripotency and the gain of acquired lineage characteristics. To determine the global molecular dynamics of differentiation, mRNA expression changes were assessed via microarray of bulk cell material, and morphological and protein expression changes were examined by immunostaining (**Fig. 1a**). To extract general, rather than cell-line specific, processes we conducted two biological replicates, starting with ESCs derived from mice with different genetic backgrounds (R1 and E14tg2a [E14] strains). We observed that in both cases, cells of the starting population abundantly expressed proteins related to the pluripotent state (**Fig. 1b**), while at the final time point of the differentiation trajectory (168 h), cells were marked primarily by neuronal stem cell marker *Sox1* and early neuronal marker *Tubb3* (**Fig. 1b** and **Supplementary Fig. S1a-b**), indicating a predominantly neuroprogenitor cell (NPC) phenotype.

**Figure 1:**
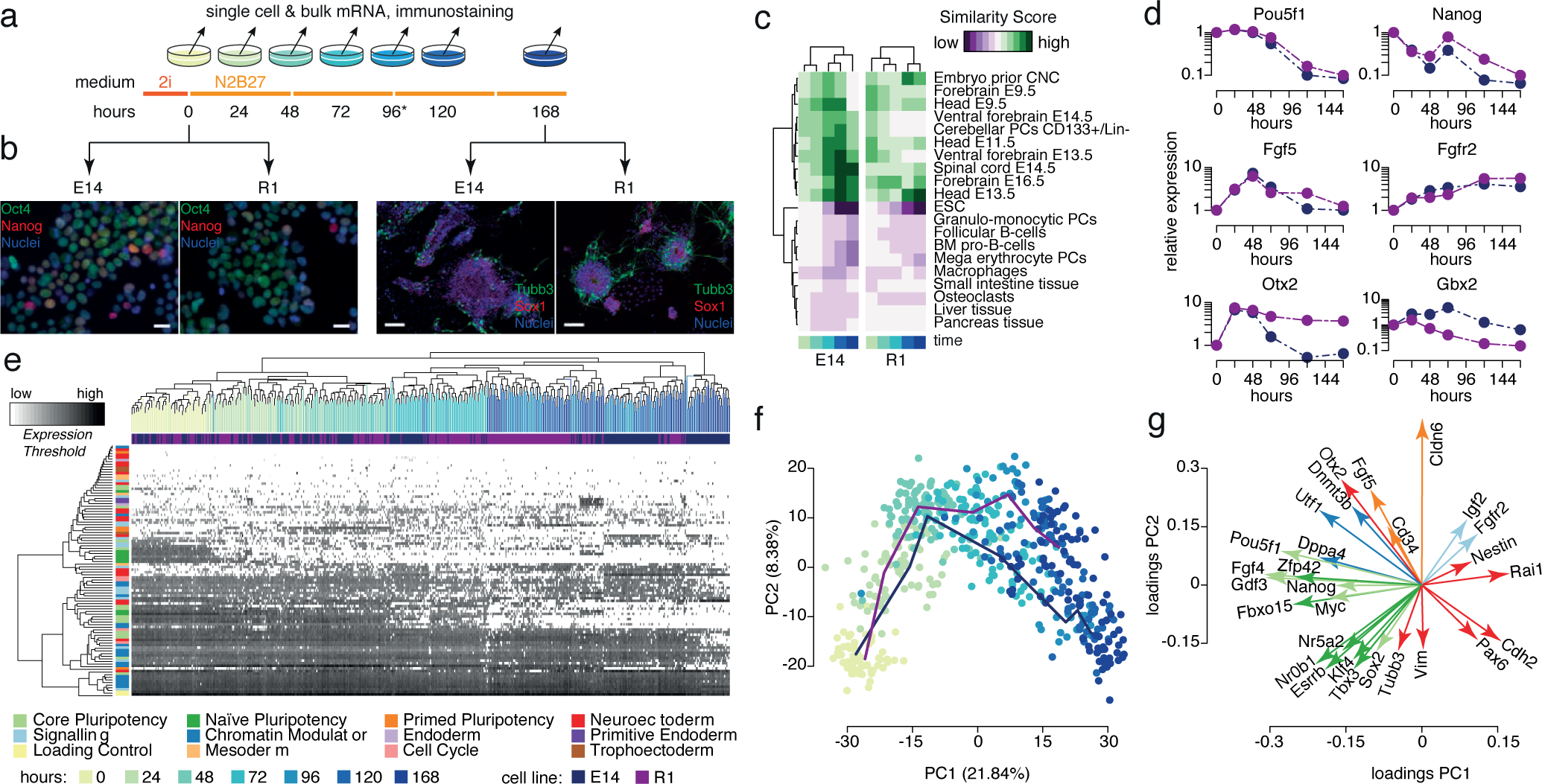
Differentiation *in vitro* recapitulates development *in vivo*. (**a**) Schematic of experimental design. (**b**) Immunostaining for pluripotency markers Oct4 and Nanog from cells at the start of the experiment (left) and neuronal markers Tubb3b and Sox1 at the end of the experiment (right). (**c**) Comparison of global gene expression profiles with a training library shows loss of pluripotency characteristics and progressive gain of neuronal characteristics. Comparisons with the 20 most similar/dissimilar lineages are shown. The full comparison is shown in **Supplementary Fig. S1**. (**d**) Loss of pluripotency markers and gain of neuronal lineage markers assessed by RT-PCR. (**e**) Single cell data shows a gradual drift from the ESC state to the NPC state. (**f**) Projection of the data onto the first 2 principal components reveals the presence of a transient intermediate state during differentiation. Solid lines show mean trajectories for each cell line. (**g**) Gene loadings for the first two principal components indicates that the intermediate state is a primed epiblast-like state. Data for the R1 cell line is given in blue, and the E14 cell line is given in purple throughout.

To better understand the dynamics of the transition from the ESC state to the NPC state we constructed a supervised machine learning classifier that compares the observed gene expression patterns to those from a training library of 161 cell-type specific gene expression profiles curated from the literature (for complete list see **Supplementary Table S1**) and produces a similarity score for each lineage based upon our previously published methodology (Lenz et al. 2013). This analysis revealed a gradual loss over time of gene expression characteristics associated with pluripotency and early development, and a sequential emergence of gene expression patterns related to the neural tube and brain development, in accordance with the appropriate mouse developmental stages (**Fig. 1c, Supplementary Fig. S2** and **Supplementary Table S2**). In particular, we observed that gene expression patterns became increasingly similar to those seen during specific stages of the head and ventral forebrain development (E9.5–E16.5), while similarity to tissues of mesodermal and endodermal origin was either consistently low or progressively reduced over time. Complementary analysis of global gene expression changes identified 1726 diferentially expressed genes throughout the time course with substantial overlap between the two cell lines (**Supplementary Fig. S1f-h**). Among those 877 consistently up-regulated genes, annotation terms for brain tissue and neuron differentiation were significantly over-represented (*p* = 8.1 × 10^−3^ and *p* = 2.9 × 10^−8^ false discovery rate [FDR] corrected, respectively), while annotations for embryonic stem cell and stem cell maintenance were enriched among the 849 down-regulated genes (*p* = 1.7 × 10^−3^ and *p* = 8.9 × 10^−3^ FDR corrected, respectively) (**Supplementary Table S3**). These results indicate the induction of appropriate, and broadly similar, differentiation programs in both cell-lines. However, subtle differences in gene expression changes between cell-lines were also apparent, indicating the initiation of slightly different developmental programs. Expression of *Otx2*, an important regulator of rostral brain development (Simeone et al. 1992), occurred only transiently during the first 48 h of differentiation in E14 cells, while expression was sustained in R1 cells (**Fig. 1d**). Concomitant with this, expression of *Gbx2*, an antagonist of rostral brain patterning during the formation of the mid/hindbrain junction (Millett et al. 1999, Broccoli et al. 1999), was subsequently induced in E14 but not in R1 cells (**Fig. 1d**). Similarly, strong expression of *Hoxa2* and *Hoxb2*, regulators of rhombomeric segmental identities in the hindbrain (Sham et al. 1993, Nonchev et al. 1996), was observed in E14 cells from 72 h onward yet only weakly in R1 cells (Supplementary Fig. S1i), suggesting a specification bias intrinsic to each cell line. Taken together these analyses indicate that differentiation *in vitro* reliably recapitulates developmental dynamics *in vivo.*

### Differentiation progresses through an intermediary metastable state

To investigate the dynamics of differentiation further we sought to monitor differentiation dynamics at the single cell level. To do so, gene expression changes for 96 pre-selected genes of interest (including regulators of pluripotency and neuronal differentiation, as well as epigenetic and cell cycle regulators, see **Supplementary Table S4**) were recorded periodically over the course of the time series within individual cells using a high-throughput RT-PCR array (**Fig. 1a,e** and **Fig. 2a**). Hierarchical clustering of this data largely captured the natural ordering by sampling time, indicating a gradual progression of cellular identities away from the ESC state towards the NPC state (**Fig. 1e**). Dimensionality reduction using principal components (PC) analysis suggested that cells do not move directly from the ESC state to the NPC state, but rather pass through a transitory intermediate state (**Fig. 1f**). Analysis of the contribution of the gene loadings to each of the first two PCs revealed that the dynamics may be decomposed into two distinct molecular processes (**Fig. 1g**): PC1 associates with the transition from the ground state of pluripotency toward the neuronal lineages (regulators of the ground state ESC identity such as *Pou5f1*, *Nanog*, *Esrrb*, *Zfp42*, *Klf4*, *Tbx3*, *Nr0b1* and *Myc* are negatively associated with this component; while genes associated with the NPC identity such as *Nestin*, *Rai1*, *Pax6* and *Cdh2* are positively associated); while PC2 associates with the process of epiblast maturation (regulators of the primed epiblast that forms the egg cylinder, such as *Otx2*, *Fgf5*, *Cd34* and *Cldn6* as well as generic epigenetic regulators such as *Utf1* and *Dnmt3b* are positively associated with this component; while characteristic neuronal genes such as *Vim* and *Tubb3* are negatively associated) (**Fig. 1g**). This analysis suggests that differentiation progresses through three biologically distinct cell states.

To further determine if this partition into three states is a strong feature of the data we conducted *k*-means clustering for 2-10 clusters and analyzed cluster qualities using the GAP statistic, a simple metric which compares the within-cluster variability present for a given clustering to that expected from appropriate randomization (Tibshirani et al. 2001), in order to identify natural clustering patterns in the data. This analysis revealed the presence of 3 robust clusters in the data and thereby confirmed that the biologically-intuitive partition of differentiation into 3 distinct phases is a natural feature of data (**Fig. 2b-d** and **Supplementary Fig. S3a**). Taken together these analyses replicate consistently-observed results from prior studies (Abranches et al. 2009, Boroviak et al. 2014, Kalkan & Smith 2014) and indicate that differentiation progresses through three phenotypically-distinct cell states: from the ground state of pluripotency to a primed epiblast-like state before the commitment to neural lineage is specifically made.

**Figure 2:**
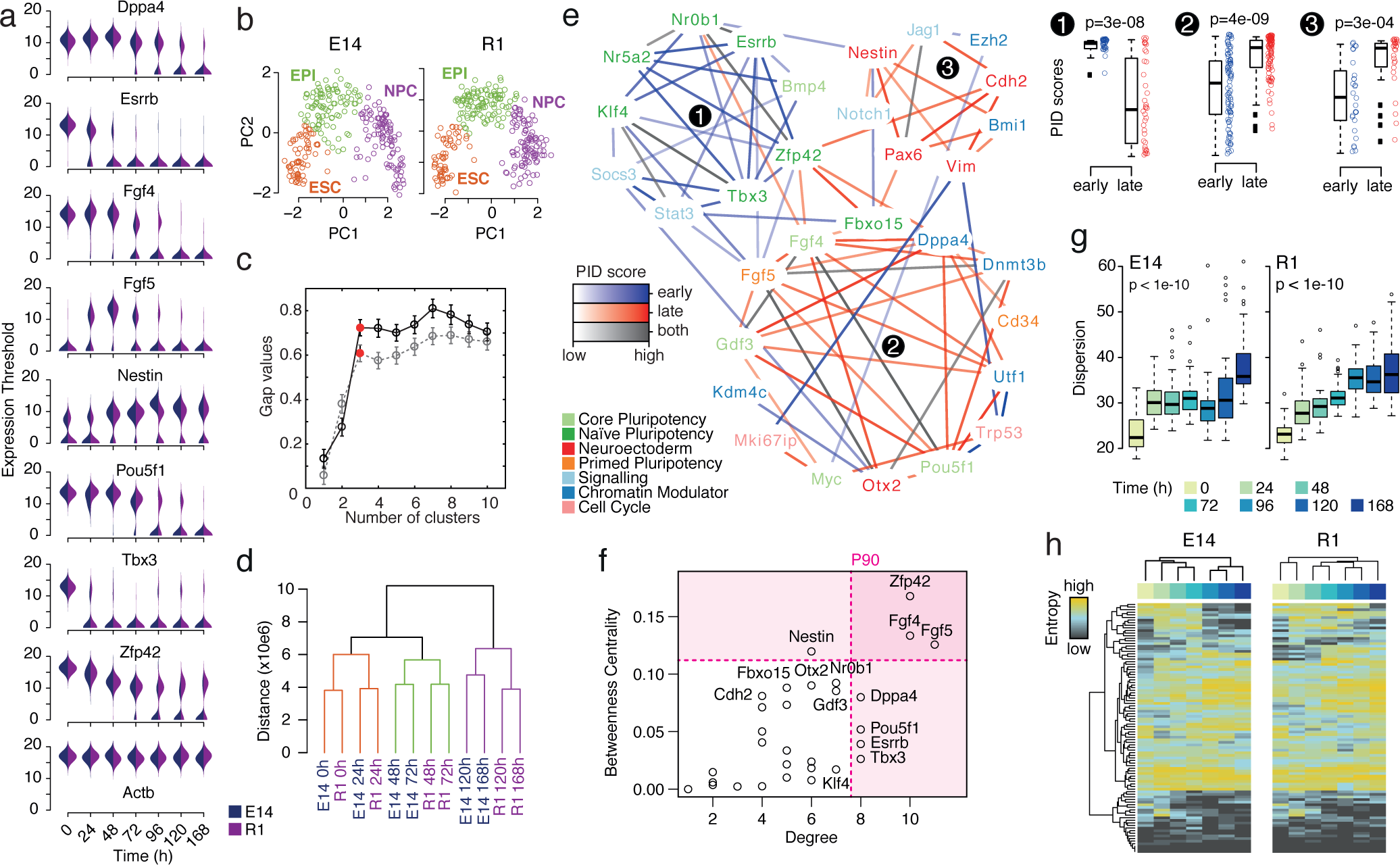
Differentiation is accompanied by regulatory network re-configurations and an increase in cell-cell variability. **(a)** Bean plots of expression changes of key genes from single cell RT-PCR data. **(b)** Single cell expression data naturally clusters into 3 distinct groups. **(c)** Assessment of cluster quality using the GAP statistic (Tibshirani et al. 2001). The most natural partition of the data is associated with the ‘elbow’ in this plot, here at three clusters highlighted in red. Error bars show standard errors. **(d)** Microarray expression data also naturally clusters into 3 distinct groups. **(e)** Regulatory network inferred from single cell data has three distinct modules that are active at different times during differentiation. Box plots to the right show the distributions of PID scores, which measure edge importance (see **Methods**), for all edges in each cluster at early and late times. Boxes show first and third quartiles about the median, whiskers extend to 1.5 times the interquartile range from the box. Data points beyond whiskers are shown as outliers above or below boxes; all the data points are shown beside the boxes. Significant changes in PID scores indicate differential expression of the module over time. p-values were obtained using a Wilcoxon rank-sum test. **(f)** Genes with high degree are important for consolidating cellular identities in each state. Genes with high betweenness centrality are important in the transition between states. **(g)** Cell-cell variability, as assessed by multivariate dispersion (see **Methods**) increases over the time-course. Boxes show first and third quartiles about the median, whiskers extend to 1.5 times the interquartile range from the box. Data points beyond whiskers are shown as outliers. **(h)** Shannon entropy, as a measure of gene expression noise, also generally increases over the time-course.

### Cell state changes are accompanied by regulatory network re-configurations

Having identified three robust cell states, we wanted to better understand the transcriptional changes that occur as cells move from one state to another, and to identify functional relationships between genes that mediate these transitions. We reasoned that if two genes are co-regulated, or if one gene regulates the other, then we would observe coordinated changes in the expression levels of these genes over time. We therefore sought to infer a putative regulatory network from the data, in order to better understand any patterns in these coordinated changes. To do so we made use of information-theoretic measures that enable the identification of non-linear statistical relationships between variables (here, genes), and are therefore substantially more powerful that traditional correlation-based network inference approaches (Villaverde et al. 2013, Mc Mahon et al.2014). In particular, we used the partial information decomposition (PID), a recently derived method to examine the statistical relationships between three or more variables that provides a more detailed description of statistical relationships than standard information-theoretic measures such as pairwise mutual information (Williams & Beer 2010, Timme et al. 2014). Our PID-based algorithm assigns a score to each potential gene-gene interaction indicating the strength of statistical association, which we interpret to be evidence of a putative functional relationship, and selects only those interactions that pass a stringent selection criterion (see (Chan et al. 2016) and **Methods** for full details). This analysis revealed a network enriched with connections between known regulators of pluripotency and neuronal differentiation (**Fig. 2e**). To dissect how regulatory interactions change over time we applied this method to different subsets of the data: to infer interactions important for the early stages of differentiation we used data from cells identified as being in the ESC and epiblast-like (EPI) states; to identify interactions important for the later stages of differentiation we used data from cells identified as being in the EPI or NPC states (individual cells were identified as being in the ESC, EPI or NPC state via k-means clustering, as described above). This analysis revealed strong clustering of edges according to their temporal importance (as colored in **Fig. 2e** and **Supplementary Fig. S3b**). To investigate this clustering further we then identified regulatory modules within the network using an unsupervised community detection algorithm that identifies modules across different scales without assuming a fixed number of modules in advance (Delvenne et al. 2010, Lambiotte et al. 2008, Schaub et al. 2012). This analysis revealed the presence of seven regulatory modules (**Supplementary Fig. S3b**), three of which displayed significant changes in activity over time (**Fig. 2e**). Genes in module 1 are primarily associated with the ground state of pluripotency (see **Supplementary Table S4** for gene annotations) and reduce substantially in expression during the early stages of differentiation. Genes in module 2 are primarily associated with the primed epiblast-like state and are generally transiently up-regulated towards the middle of the time-series and down-regulated from approximately 72 h onward. Genes in module 3 are primarily associated with neuroectoderm differentiation, and generally increase in expression throughout the time-course. While most genes within each of these 3 modules primarily displays strong intra-module connectivity (that is, they connect strongly to other members of the same module but weakly to members of different modules), some genes such as *Zfp42*, *Fgf5*, *Fgf4* and *Nestin* also showed high inter-module connectivity (as assessed by betweenness centrality, a simple measure of node importance, see (Newman 2010) and **Methods**), suggesting a potential role for these genes in coordinating the transitions between states (**Fig. 2f**). In contrast, those genes that form the hubs of their respective modules, such as *Dppa4* and *Pou5f1* (**Fig. 2f**) may be involved in the maintenance or consolidation of one particular cell state. Collectively, these results reaffirm that the early stages of differentiation progress through two distinct pluripotent states and indicate that coordinated changes in regulatory network structure accompany these cell state changes.

### Gene expression noise increases during differentiation

Once we had identified these three states, we sought to better understand the dynamics of cellular transitions between states. We reasoned that if cells pass from one state to another in a coordinated deterministic way then the initial cell-cell variability present in the population would propagate with time, and therefore cell-cell variability would remain approximately constant or reduce through the time-series. Alternatively, if cells were to progress in an uncoordinated, stochastic way from one state to another, then cell-cell variability would increase over time. To investigate this we estimated the total dispersion within the population from our single cell expression data. Dispersion is a multivariate measure of cell-cell variability that takes into account the variability of each gene as well as the patterns of covariance between genes (see **Methods**). This analysis revealed a significant increase in cell-cell variability over time (**Fig. 2g**). To investigate this increase further, we also estimated the Shannon entropy of expression for each gene at each time, which may be thought of as a simple measure of expression disorder (MacArthur & Lemischka 2013). This analysis also revealed a general increase in gene expression noise as differentiation progresses (**Fig. 2h**). Taken together, these results suggest that while all cells are exposed to the same differentiation cues, cellular differentiation in response to these cues progresses in an uncoordinated and apparently stochastic way.

### Differentiation is a stochastic process with memory

Taken together, our statistical analysis confirmed the widely-accepted model that differentiation progresses through three functional cell states: from the initial ESC state, to a primed epiblast-like (EPI) state, and then on to the final NPC state. However, the increase in cell-cell variability we observed also indicated that cells do not synchronize their transitions through these states. Rather it appeared that individual cells transition through these states in an uncoordinated, stochastic manner. We reasoned that this inherent stochasticity might be important, yet the mechanisms by which it is regulated were not clear. To investigate further we sought to construct a series of mathematical models to explore this process (for details, see **Box 1**). In our first, most basic model, we assumed that cells are initially held in the naïve pluripotent state when cultured 2i conditions, yet once these extrinsic constraints are released cells progress stochastically from one state to the next at constant average rates (see schematic in **Figure 3a** and details in **Box 1**). We found that this first model does not describe the data well (**Figure 3a**), since it allows cells to transition quickly through the ESC, EPI and NPC states, yet we observed that the first pioneer neurons emerge *in vitro* only after 72-96 h (**Supplementary Fig. S1c**), corresponding to the same phenomenon in mouse corticogenesis from E8.5 onward (Stainier & Gilbert 1990). Thus, while the majority of cells accumulate in the EPI state around 72 h in experiment, the model cannot account for this accumulation. This suggests that individual cells within each state are not interchangeable with respect to their differentiation potential, but rather are distinguished from one another with respect to some hidden (that is, unmeasured) variables. To account for these hidden states we next sought to model the dynamics using a hidden Markov model, a stochastic modeling framework that accounts for the effects of unobserved variables on the observable dynamics (Rabiner 1989). In our revised model we allowed the ESC, EPI and NPC ‘macrostates’ to conceal a chain of *N* ‘microstates’, which the cells transit through stochastically in sequence at a constant average rate (**see Box 1**). We found that this simple revision describes the experimental data remarkably well (**Fig. 3b**). Model fitting indicated the presence of 8 hidden microstates within the observed ESC state for both cell-lines, and 11 (12) microstates within the observed EPI state for R1 cells (E14 cells respectively). The expected transition time between microstates was 5.3 (4.8) h for R1 cells (E14 cells respectively), giving a mean residence times of 42.6 (40.8) h in the ESC state, and 63.9 (56.1) h in the EPI state for R1 cells (E14 cells respectively) (**Fig. 3c**). It is of note that the inferred transition times between microstates are significantly shorter than the cell cycle time, which is approximately 15 h for these cells in both 2i and N2B27 media (**Fig. 3d**), while the inferred transition times between macrostates are significantly longer than the cell cycle time. This suggests that the dynamics are not primarily driven by cell division events, but rather by some as yet unidentified molecular processes. However, the surprising consistency in the fitted model parameter values between the two cell lines gives reliability to these model estimates and suggests that both lines are undergoing a common dynamical process, with the same number of intermediate transition states, despite their slight molecular differences. This analysis therefore suggests that stem cell differentiation along the neuronal lineage is an inherently discrete, yet strongly canalized, stochastic process.

**Figure 3:**
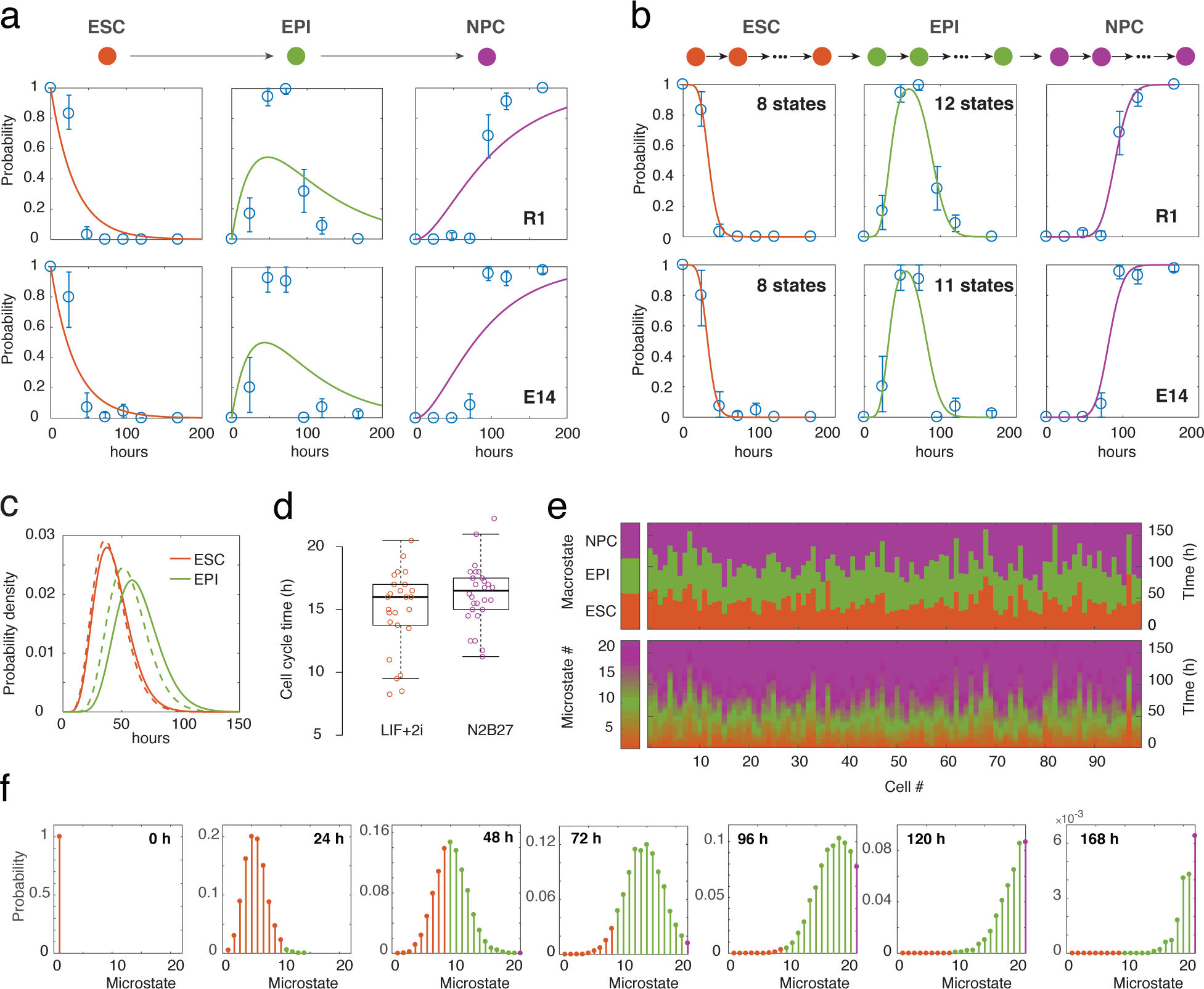
Data fitting to a hidden Markov model reveals the presence of cellular microstates. (**a**) Fit of data to Eqs. (1) – (3). This memoryless stochastic process does not describe the data well. (**b**) Fit of data to Eq. (10). Data is well described by this stochastic process with memory. (**c**) Wait times in the ESC and EPI states. Full lines show E14 data, dotted lines show the R1 data. (**d**) Cell cycle times in LIF + 2i and N2B27 media are significantly longer than microstate residence time. (**e**) Illustrative simulation of 100 cells according to our hidden Markov model. Parameters are taken from the R1 model fit. (**f**) The resulting probability density function over the microstates. Throughout this figure orange represents the ESC state; green represents the EPI state; and purple represents the NPC state.

##### Box 1: Mathematical models

Let *p_A_*(*t*), *P_B_*(*t*) and *p_C_*(*t*) be the probabilities that a randomly selected cell is in the ESC, EPI or NPC state respectively at experimental time *t*. Assuming that all cells within a given state behave in the same way and transitions between states occur independently at constant average rates, these dynamics are described by the following set of equations:

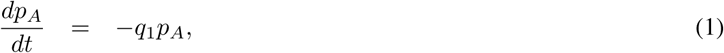

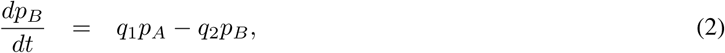

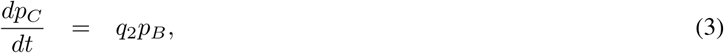

where *q*_1_ and *q*_2_ are transition probabilities per unit time, and we assume that *p_A_*(0) = 1 and *p_B_*(0) = *p_C_*(0) = 0 (i.e. all cells start in the ESC state). This model, which assumes that cells within each observable state are homogeneous with respect to their differentiation potential, does not describe the data well (see Fig. 3a). This suggests that individual cells within each observable state are not interchangeable, but rather are distinguished from one another with respect to some hidden variables. To account for these differences we modified this model to allow each observable ‘macrostate’ to conceal a number of hidden ‘microstates’. Let *p_n_* be the probability that a cell is at microstate *n* at time *t*. For simplicity we assume that cells transition from one microstate to the next on average at the same rate. In this case, the dynamics of the hidden Markov process are given by

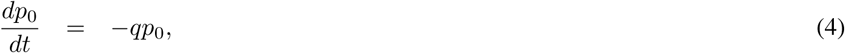

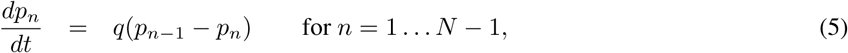

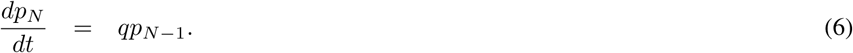

where *q* is the transition probability per unit time, with *p_n_*(0) = *δ*_*n*0_ (i.e. all cells start in the first microstate). This model is simply a homogeneous Poisson process, and may be solved exactly (Van Kampen 2007) to give

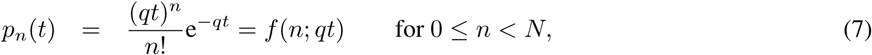

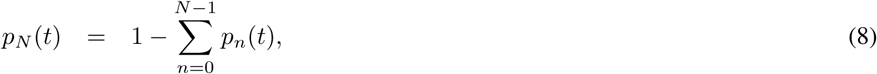

where *f*(*n*; *q_t_*) is the Poisson probability density function. Assuming that microstates 0,1,2,…, *n_A_* identify with the ESC state, microstates *n_A_* + 1, *n_A_* + 2,…, *n_B_* identify with the EPI state, and microstates *n_B_* + 1, *n_B_* + 2,…, *N* identify with the NPC state, the observed probabilities,

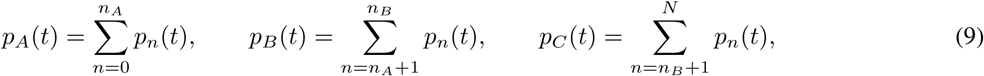

may also easily be found,

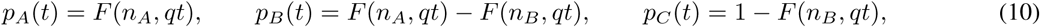

where *F*(*n*; *qt*) is the Poisson cumulative distribution function. The dynamics of this model are illustrated in **Fig. 3e-f**.

### Hematopoietic stem cell differentiation follows similar dynamics

Taken together, our analysis suggests that ESC differentiation is a robust, yet inherently discrete stochastic, process. However, although we saw a strong consistency between the two cell lines it was not clear if the dynamics we observed were particular to ESCs or had a more general relevance. To explore our modeling framework further we next sought to determine if it could be used to describe differentiation in an unrelated biological context, using a different molecular profiling technology. To do so, we considered the dynamics of surface marker expression, as assessed by FACS, in long-term repopulating hematopoietic stem cell (LT-HSC) as they differentiated along the myeloid lineage. Lin^−^Sca-1^+^c-Kit^+^CD41^−^CD48^−^CD150^+^CD34^−^ LT-HSCs were isolated from 8 week-old C57BL6 mice and cultured in conditions that promote myeloid differentiation, according to a well-established protocol (Rieger et al. 2009). Cells were sampled at 3, 5, 7, 10, and 14 days after induction, and each cell was classified as being in a stem, progenitor or committed state, depending on its expression of surface markers Sca-1 and c-Kit (see **Fig. 4a** and **Methods**). We found that our hidden Markov model also described these dynamics remarkably well (**Fig. 4b**), suggesting that it provides a general framework to understand mammalian stem cell dynamics. In this case model fitting indicated the presence of 6 hidden microstates within the observed LT-HSC state, and 3 hidden microstates within the progenitor state, with an expected transition time between microstates of 26.7 h. These model parameters differ substantially from those observed during ESC differentiation. Thus, while differentiation of LT-HSCs may be described by the same general framework as ESC differentiation, the qualitative characteristics are quite different, suggesting that hematopoietic stem cell differentiation is driven by distinct underlying molecular processes.

**Figure 4:**
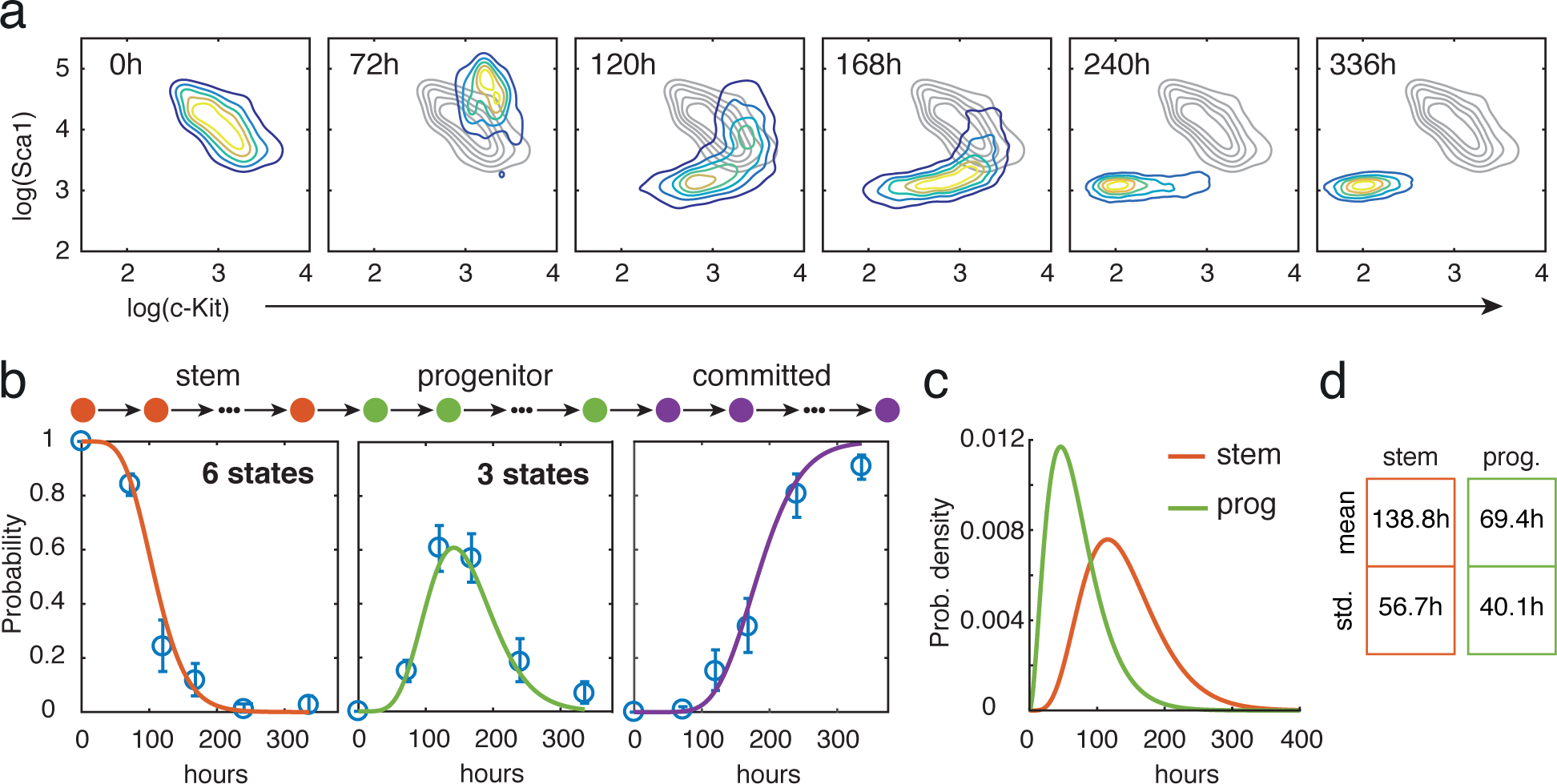
Hidden markov model of LT-HSC differentiation. (**a**) Expression changes in Seal and c-Kit characterizes the dynamics of differentiating LT-HSCs. (**b**) Fit of data to Eq. (10). Data is well-described by a stochastic process with memory. (**c**) Wait time distributions in the LT-HSC and progenitor states. (**d**) Mean and variance of the residence times in the LT-HSC (green) and progenitor (orange) states. Wait-times are substantially more variable for LT-HSC differentiation than for ESC differentiation, indicating an inherently more stochastic process.

## Conclusions

Recent years have seen remarkable advances in high-throughput single cell profiling technologies (Stegle et al. 2015, Grün & van Oudenaarden 2015, Shapiro et al. 2013, Hoppe et al. 2014, Kolodziejczyk, Kim, Svensson, Marioni & Teichmann 2015, Moignard & Göttgens 2016). To better understand the data that these new and emerging methods produce, there is now a need for modeling and analysis methods that sift functional cell-cell variability from measurement noise, and identify distinct cellular identities from highly heterogeneous data. These issues are particularly apparent when considering time-course data and a number of computational tools have accordingly recently been developed to explore cell fate trajectories from single cell time-course data, for example using pseudotemporal ordering (Trapnell et al. 2014, Bendall et al. 2014, Setty et al. 2016, Haghverdi et al. 2016). These computational models are typically based on the assumption that cells progress continuously through measurable cell states and so implicitly assume that underlying molecular stochasticity is buffered to the extent that the continuum approximation is appropriate. However, it has been observed that combinatorial fluctuations in key lineage-specifying factors (Chambers et al. 2007, Toyooka et al. 2008, Hayashi et al. 2008, MacArthur & Lemischka 2013, Abranches et al. 2014) are important for stem cell fate specification, and it has been argued that cell fate commitment is accordingly a fundamentally discrete stochastic process (Moris et al. 2016, Pina et al. 2012).

Here, we have outlined an alternative modeling framework that infers the presence of discrete hidden cell states from limited expression data and have used this framework to dissect the dynamics of neuronal differentiation of mouse ESCs *in vitro.* In accordance with previous observations we find that differentiation progresses through two functionally distinct pluripotent cell states: a naïve pluripotent state representative of the transient ESC state *in vivo* and a primed-pluripotent state, representative of the post-implantation epiblast *in vivo* (Abranches et al. 2009, Boroviak et al. 2014). However, our analysis also indicates that these observed states conceal a multitude of functionally-similar hidden cell states. By analogy with statistical mechanics (MacArthur & Lemischka 2013, Garcia-Ojalvo & Arias 2012, Moris et al. 2016), we refer to the observable functional cell states as cellular *macrostates*, and the variety of molecular configurations associated with each functional macrostate as cellular *microstates*. In our framework the microscopic dynamics are given by a homogeneous Poisson process (Van Kampen 2007) in which the number of system states is a hidden variable. Since the probability that a cell will transition to the next microstate per unit of time is independent of how long it has spent in its current microstate this underlying stochastic process is Markovian (or memoryless). However, transitions between macrostates are not Markovian: the probability that a cell will transition to the next macrostate depends on how long it has already spent in the current macrostate. Thus, in our formalism the macroscopic dynamics, which describe transitions between functional cell types, are, formally, a stochastic process with memory (see **Fig. 5** for a schematic, and **Supplementary materials** for more details). During differentiation this memory is important since it allows individual cells to keep a record of their progress, and provides a simple epigenetic mechanism by which cells can consolidate a particular functional identity before progressing onto the next. In our view, the interplay between microstates and macrostates is therefore fundamental to the process of differentiation.

**Figure 5:**
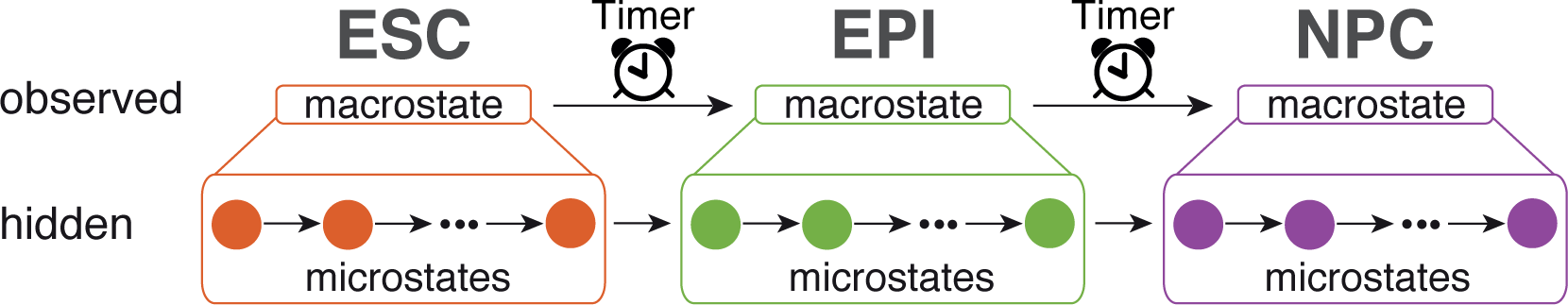
Schematic of model framework. Cells transition at a constant rate through a chain of hidden microstates which are not directly observed but rather group together into observable macrostates. While the underlying dynamics are Markovian, the observable dynamics are non-Markovian, and may therefore be thought of as a stochastic process with memory.

For example, the number of microstates in the differentiation trajectory has an important role in regulating its output. In the case of differentiation of mouse ESCs along the neuronal lineage we estimate that there are 20-21 states in the chain (**Fig. 3b**). Thus, while each transition from one microstate to the next is inherently stochastic, a large number of these transitions must occur in order for the cell to differentiate fully. In stochastic analysis it is well-known that the output of such a chain of stochastic events becomes less variable as the length of the chain increases, a result that is known as the law of large numbers (Gardiner 1985). In our model, this means that the length of time it takes for an individual cell to complete the differentiation trajectory becomes less variable as the number of microstates on the trajectory increases. The large number of microstates we estimate in the chain between the ESC and the NPC states therefore serves to regularize an inherently stochastic process, and ensure that differentiation occurs in a reliable and reproducible way. By contrast, since we estimate that there are only 9 hidden states in the transition from the hematopoietic stem cell state to the committed myeloid state, the output of this trajectory is correspondingly more variable (compare the spread of the wait time distributions in **Fig. 2c** with those in **Fig. 4b**). For this reason hematopoietic stem cell (HSC) differentiation appears to be more stochastic than ESC differentiation.

This observation may reflect differences in the functional roles that ESCs and HSCs play. All stem cells must balance adaptability (the ability to respond quickly and appropriately to environmental changes) with robustness (the ability to reliably differentiate when required). However, the relative importance of these opposing constraints in regulating cell behavior inevitably depends on the particular stem cell type. In their natural context, ESCs are only present for a short period of time during early development, and must reliably initiate the production of a wide variety of tissues during this short time. In this developmental context, it is essential that differentiation occurs in a robust and reproducible way: excessive stochasticity must therefore be constrained and the insertion of a large number of transient states in the differentiation trajectory is a simple, and parsimonious, way to do this. By contrast, hematopoietic stem cells reside throughout life in the adult and are responsible for maintaining healthy maintenance of the blood lineages, and rapid replacement of tissue when needed. In this primarily homeostatic context, adaptability is of greater relative importance than robustness. Here, stochasticity may be an advantage since it simultaneously allows the maintenance of healthy tissue balance and the ability to remain primed, or poised, to respond to environmental changes as needed. In summary, our analysis indicates stem cell differentiation is well-described by a simple stochastic process, and is amenable to analysis using the tools of statistical mechanics. We anticipate that the most exciting future advances in stem cell science will combine new experimental techniques with further theoretical developments in the physics of living matter.

## Methods

### Experimental Procedures

#### Routine cell culture media

Pluripotent mouse embryonic stem cells (mESC) were cultivated in Dulbecco’s Modified Eagle Medium (DMEM; life technologies, Paisley, UK, #31053-028) with 1% Penicillin/Streptomycin (PAA, Yeovil, UK, #P11-10) that was further supplemented with 15% KnockOut serum replacement, 1x MEM non-essential amino acids, 1x GlutaMax (all from life technologies, Paisley, UK, #10828-010, #11140-050 and #35050-038), 50 *μ*M 2-mercaptoethanol (Sigma Aldrich, Gillingham, UK, #M6250). Leukaemia inhibitory factor (LIF), produced in house, was added at a saturating dilution of 1:1000. Cells were seeded on 0.1% gelatine (Sigma-Aldrich, Gillingham, UK, Cat. No. G1890) coated tissue culture plates pre-seeded with *γ*-irradiated MEF for routine culture. Throughout four subsequent passages prior to the start of the experiment, cells were cultivated in 0.1% gelatine coated tissue culture plates without additional MEF, and medium was additionally supplemented with a combination of 1 *μ*M PD0325901 (Tocris bioscience, #4192) and 10 *μ*M CHIR99021 (Reagents Direct, #27-H76). Cells were maintained at 37°C and 5% CO_2_ and routinely passaged every other day using Trypsin/EDTA (PAA, Yeovil, UK, #L11-003). Medium was replaced on a daily basis.

##### Neuronal differentiation

Neuronal differentiation medium (N2B27) was prepared according as previously described (Ying et al. 2003) and contained a mixture of Neurobasal and DMEM/F12 media, supplemented with B27 and N2 supplements (Thermo Fisher, Cat.No. 12348017, 21041025, 17504044 and 17502048).

##### Myeloid differentiation of hematopoietic stem cells

LT-HSCs (Lin^−^Sca-1^+^c-Kit^+^CD41^−^CD48^−^CD150^+^CD34^−^ cells) were isolated from 8 week-old C57BL6 mice and differentiated as previously described (Rieger et al. 2009). Cells were cultured in 96-well flat bottom plates in StemSpan SFEM (StemCell Technologies) in the presence of stem cell factor (SCF, 100 ng/ml, PeproTech), IL-3 (10 ng/ml, PeproTech), IL-6 (10 ng/ml, PeproTech), M-CSF (50 ng/ml, PeproTech) and 10^−5^M 2-mercapto ethanol (2-ME, Invitrogen) at 37°C, 5% CO2. After 3, 5, 7, 10, and 14 days of culture, cells were collected and the expression of Sca-1, c-Kit were quantified by FACS. Data was normalised using the corresponding isotype control to account for technical variation in day-to-day fluorescence acquisition. Background signal intensities for each fluorescence channel were estimated from the modes of the isotype controls and a correction factor was calculated based on the shift required to center these modes around the same value. Intensity values of the stained specimens were then adjusted by the correction factor of their corresponding isotype control. Cells were classified as being in either a stem, progenitor or committed state using *k*-means clustering with 3 clusters and associating the Sca-1 high/c-Kit high cluster with the stem cell state; the Sca-1 low/c-Kit high cluster with the progenitor state; and the Sca-1 low/c-Kit low cluster with the committed cell state.

##### Isolation of mRNA

Total mRNA was isolated from cell lysates according to manufacturer’s instructions using the AllPrep DNA/RNA Mini Kit (Quiagen, Crawley, UK, Cat.No. 80204).

##### Global Gene Expression Microarrays

For global gene expression, total mRNA isolated from ensemble cells was processed and hybridized to MouseWG-6 v2.0 Expression BeadChip mircoarrays by CGS genomics, Cambridge, UK. Differentially expressed genes (DEG) were identified based on their relative expression changes throughout the entire differentiation time course, denoted as cumulative relative expression (CRE). A gene was considered a DEG when it’s CRE surpassed a threshold of 3 times the interquartile range above/below the 75 percentile (25 percentile) based on the entirety of CREs.

##### Single cell gene expression arrays

Individual cells were sorted using a BD FACS Aria II flow cytometer into 96-well round bottom multi-well plates (both Becton-Dickinson, Oxford, UK). Single cells were de-posited directly into 5 *μ*l of reaction mix containing reagents for cell lysis, reverse transcription, as well as the polymerase and reaction buffers for RT-PCR. The reaction mix consisted of 0.1 *μ*l Superscript III RT/Platinum Taq Mix, 0.05 *μ*l Ambion’s SUPERase12-In, 1.85 *μ*l DEPC-treated water (all part of CellsDirect One-Step qRT-PCR Kit, life technologies, Paisley, UK, Cat. No. 11753) and 0.0125 *μ*l of 96 different TaqMan assays (probe IDs included in supplementary table) for multiplex pre-amplification. The reverse transcription and pre-amplification was performed on a Veriti thermal cycler (life technologies, Paisley, UK) with the following temperature cycles: 15 min, 50°C; 2 min, 95°C followed by 22 cycles of 15 s at 95°C alternating with 1 min at 60°C. Thus, pre-amplified cDNA was diluted with 20 *μ*l of DEPC-treated water and stored at −80°C until further processing. Readout was performed using Fluidigm 96x96 Dynamic Array in combination with the Biomark HD system (both Fluidigm, San Francisco, USA) according to manufacturers instructions.

##### Immunofluorescence staining

Cells were fixed for 20min at room temperature (RT) using 4% Paraformaldehyde (Sigma-Aldrich, Gillingham, UK, #P6148) in PBS-/- (PAA, Yeovil, UK, #H15-002) and washed three times with PBS-/-. Intracellular epitopes were made accessible by permeabilisation of the cell and nuclear membranes using a 0.2% Triton-X-100 (Sigma-Aldrich, Gillingham, UK, #X100) solution in PBS-/-for 10min at RT. Unspecific binding sites were blocked for 45min at RT with 0.1% Triton-X-100 and 10% fetal bovine serum (life technologies, Paisley, UK, #10270106) in PBS-/-, washed three more times before re-suspension in blocking buffer and either primary antibody or matching isotype controls and incubation over night at 4ÂžC under slow, continuous agitation. Cells were subsequently washed three times using blocking solution and re-suspended in blocking solution and secondary antibodies for incubation under continuous agitation for 1h at RT. Samples were washed three times in blocking solution and nuclei were stained at RT for 10min using 4’,6- diamidino-2-phenylindole (DAPI; Sigma-Aldrich, Gillingham, UK, #D9542) at a concentration of 10 *μ*g/m. Cells were imaged following a final wash in PBS-/- using a Carl Zeiss AxioVert 200 or a Nikon Ti Eclipse microscope.

##### Cell cycle time analysis

Bright field images of cells grown at 37ÂřC and 5% CO2 in either in 2i+LIF culture medium or N2B27 medium were taken in 15 min intervals using an Eclipse-Ti microscope and NIS elements v4.3 software (both Nikon, Kingston Upon Thames, UK). Cell cycle time was measured manually by tracking the number of frames between two subsequent cell division events.

#### Analysis Procedures

##### Machine-learning of cell identities

To determine how the expression patterns of the cells in our time-course related to known tissues and cell types, we collated a database of 161 tissue/cell type specific expression patterns (**Supplementary Table S1**). Raw data sets were downloaded from the Gene Expression Omnibus (GEO, http://www.ncbi.nlm.nih.gov/geo/) database and pre-processed as a single set using the robust multi-array average (RMA) normalization method in the Affymetrix Power Tools software (http://www.affymetrix.com/estore/partners_programs/programs/developer/tools/powertools.affx). The annotation of samples into tissue/cell types was performed manually based on the experimental descriptions in the GEO database. Our experimental data collected at 24h, 48h, 72h, 120h, and 168h from both cell lines (E14 and R1) were compared to the undifferentiated (0h) samples of the respective cell line and expression differences were projected onto the training set as described in (Lenz et al. 2013). Briefly, for each comparison of time points, two gene sets consisting of the top 5% of upregulated genes and top 5% of downregulated genes were defined, and their expression values in each of the 161 tissue/cell type specific expression patterns were compared using a Wilcoxon rank sum test. This resulted in 161 tissue/cell type specific scores per time point for each cell line (signed log10 p-values of Wilcoxon test), which summarize the similarity of the observed gene expression pattern with each of the 161 tissue/cell line samples we collated. Overall these evolving scores describe the differentiation dynamics in a genome-wide expression space with physiologically relevant signatures.

##### Clustering and dimensionality reduction

All clustering and dimensionality reduction was performed in R and Matlab using standard routines. We found that a more robust clustering was obtained from the single cell data by taking a binary representation of the data (i.e. retaining only information on whether each gene is expressed or not) and performing PCA, retaining the first 2 components, prior to clustering. PCA is a well-established method for data de-noising (Friedman et al. 2001) and discretization of gene expression data has been shown to improve the robustness of subsequent analysis algorithms (Tuna & Niranjan 2010). Here, these de-noising steps make the subsequent analyses more stable but do not affect any of the conclusions of the paper. The changes in the proportions of cells in each macrostate over time were determined by calculating the fraction of cells in each cluster at each time point. Confidence intervals on proportions were obtained by Bootstrap resampling.

##### Regulatory network inference

Normalized single-cell data for each gene were discretized independently using the Bayesian Blocks algorithm, a method designed to find an optimal binning for a set of values without enforcing uniform bin width (Scargle et al. 2013, Vanderplas et al. 2012). Data from both cell types (R1 and E14) were combined for this discretization step. There were 22 genes with no detected expression in greater than 80% of cells; data from these genes were removed, leaving 74 genes for all subsequent analyses.

To infer statistical dependencies between genes from the time-series data we developed an information-theoretic network inference algorithm. Many network inference algorithms exist that use the mutual information between pairs of variables as a measure of statistical dependency (Mc Mahon et al. 2014, Meyer et al. 2008, Villaverde et al. 2013). Here, we adapted these methods to calculate a score between pairs of genes that takes into account the context of the wider network, by considering the multivariate relationships of each pair of genes with every other gene in the network. This method highlights the strongest relationships for each gene, rather than simply the strongest relationships within the whole network. We find that this methods performs better than or comparably to existing information theoretic based inference methods. Full details of this algorithm, along with bench-marking against alternative methods, may be found in an accompanying paper (Chan et al. 2016). Briefly, we make use of the partial information decomposition (PID) (Timme et al. 2014) to calculate a set of multivariate information measures that encode the statistical relationships between triplets of genes, by decomposing mutual information into synergistic, redundant and unique contributions. Specifically, if we consider the information provided by a set of genes, e.g. *A* = {*X*, *Y*}, about another target variable, e.g. *Z*, the mutual information *I (*X*, *Y*;*Z*)* between the set *A* and *Z* is equal to the sum of four partial information terms,

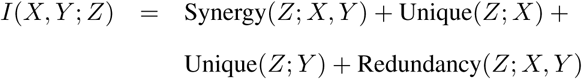

The mutual information between a single gene (*X*, say) in *A* and the target comprises a unique and redundant contribution,

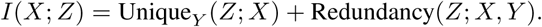

For any pair of genes, *X* and *Z*, this mutual information, *I*(*X*;*Z*), is constant regardless of the choice of the third variable, *Y*, but the unique contribution to this information varies with *Y*. Higher ratios of unique information to mutual information indicate a stronger dependency between *X* and *Z* (Chan et al. 2016). Our inference algorithm defines a measure *u_X,Z_*, based on these ratios, which we call the proportional unique contribution,

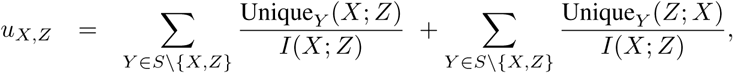

and uses this metric to assess the strength of the relationship between the pair of genes *X* and *Z*, in the context of all the other genes in the network, *Y* ∈ *S*\{*X,Z*} (where *S* is the complete set of genes). These proportional unique contributions are then used to calculate a confidence score *c*, which we call the PID score, between each pair of genes,

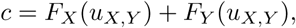

where *F_X_*(·) is a cumulative distribution function estimated using all the proportional unique contribution scores involving gene *X*. The PID scores are then used as edge weights in the (un-directed) inferred network. Edges were retained in the network if they were in the top 5 % of PID scores.

To identify molecular regulatory mechanisms active at different stages of differentiation we inferred networks from the early part of the time-course (using expression patterns from cells identified as being in the ESC or EPI states) and from the late part of the time-course (using expression patterns from cells identified as being in the EPI or NPC state).

##### Identification of modules in regulatory networks

In order to identify modules within the inferred networks that show coordinated changes in gene expression, we used a community detection method based on the evolution of a Markov process on a network, as described previously (Delvenne et al. 2010, Lambiotte et al. 2008, Schaub et al. 2012). We scanned for stable partitions at 200 Markov times from 10^−2^ to 10^2^, and selected as stable partitions those in which the number of modules remained constant for at least 10 time points, and that corresponded to a minimum in the variation of information.

##### Network analysis

Let *A_ij_* = *A_ji_* be the adjacency matrix for the network *G*. The degree of node *i* is given by Σ*_j_ A_ij_*. The betweenness centrality of node *i* is given by Σ_*j*≠*i*≠*k*_ *σ_jk_*(*i*)/*σ_jk_* where *σ_jk_* is total number of shortest paths between nodes *j* and *k* and *σ_jk_*(*i*) is the total number of shortest paths from nodes *j* to *k* that pass through node *i* (Newman 2010).

##### Estimation of dispersion and entropy

To the *i*th cell in the population we associate a gene expression vector *G*_i_ = (*g_i_*_1_,*g_i_*_2_,…, *g_i_*_96_) € ℝ^96^, which records its expression status with respect to the 96 genes we measured. Assuming that there are *n* cells in the population, the mediancentre is that point *M* = (*m_1_*, *m*_2_,…, *m*_96_) ∊ ℝ^96^ such that 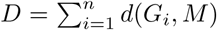 is minimum, where 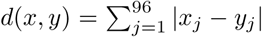 is the *L*_1_-distance. The mediancentre is a multivariate generalization of the univariate median (Gower 1974). The dispersion of each cell is its distance to mediancentre *d*(*G_i_*, *M*), and the dispersion of the population is the minimized value of *D*. The dispersion is a simple statistic that can be used in hypothesis testing to compare the multivariate variability in different populations.

To estimate gene expression entropy, normalized single-cell data for each gene were discretized independently using the Bayesian Blocks algorithm, a method designed to find an optimal binning for a set of values without enforcing uniform bin width (Scargle et al. 2013, Vanderplas et al. 2012). The Shannon entropy, *H* = −Σ*_i_ P_i_* log_2_ *P_i_*, where *P_i_* is the probability of observing gene expression in bin *i*, was then calculated directly.

##### Model fitting

Model parameters were estimated by minimizing the residual sum of squares between the data and the model. Since the model has both integer and real parameters optimization was performed using a pattern search algorithm, implemented in Matlab as part of the Global Optimization Toolbox. Models with a large number of microstates generally fitted the data better than those with a small number of microstates, since they effectively introduce more parameters into the model. To avoid over-fitting we therefore penalizing models with large numbers of microstates. Thus, we solved

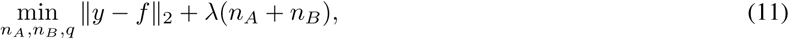

where *y* is the data and *f* is the model. The regularization parameter λ was selected using the L-curve method (Friedman et al. 2001).

## Acknowledgments

We thank R. Jewell and C. MacGuire for FACS support and Neil Smyth for the provision of LIF. M.L. was supported by the Ministry for Innovation, Science and Research of German Federal State of North Rhine-Westphalia, Germany, and the Dutch Province of Limburg, The Netherlands. This research was funded by the Biotechnology and Biological Sciences Research Council, United Kingdom, grant number BB/L000512.

## Author Contributions

P.S.S., F-J.M. and B.D.M. designed experiment; P.S.S. and R.C.S. conducted experiments; F.A. conducted single cell gene expression arrays; M.L., A.S. performed the machine learning; A.B., T.E.C. and M.P.H.S. conduced the network analysis; P.S.S., B.D.M., C.P.P and S.D.H. developed the mathematical models; P.S.S. and B.D.M. wrote the manuscript with input from all other authors.

## 7 Supplementary Materials

### Macroscopic dynamics

To describe the macroscopic dynamics we introduce the probability densities *ρ_A_*(*t*, *τ*), *ρ_B_* (*t*, *τ*), and *ρ_C_* (*t*, *τ*), where *τ* is a cell-intrinsic variable that records the length of time that an individual cell has spent in each macrostate. The observed proportion of cells in each state at experimental time *t* may then be obtained by integrating over these internal times. Thus,

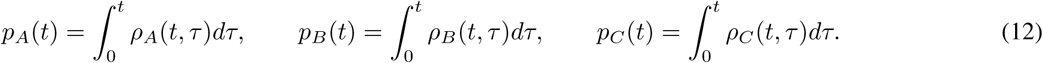

The dynamics of *ρ_A_*, *ρ_B_* and *ρ_C_* are given by the following set of partial differential equations,

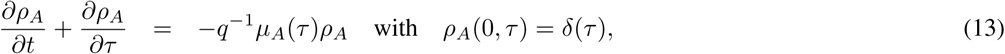

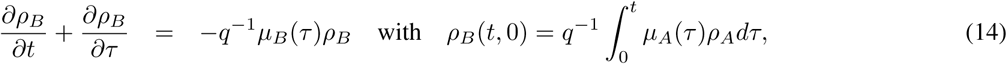

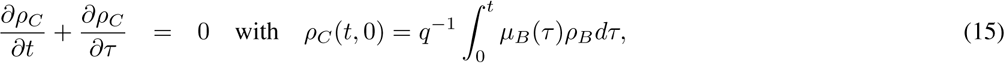

where *μ_A_*(*τ*) and *μ_B_*(*τ*) are the cumulative distribution functions for the wait times in the ESC and EPI macrostates respectively. In this case, since microscopic dynamics are given by a homogeneous Poisson process, the wait times in the ESC and EPI states are Erlang distributed (Forbes et al. 2011) and

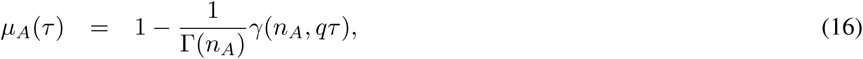

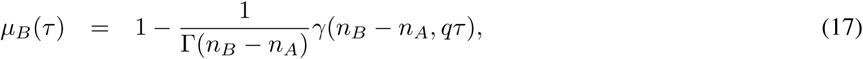

where Γ is the Gamma function and *γ* is the incomplete Gamma function. The terms on the left hand sides of Eqs. (13)–(15) account for cellular aging in each of the macrostates, while the right hand sides and boundary conditions account for transitions between macrostates. In the case that *n_A_* = 1, and *n_B_* =2, the microstates and macrostates are coincident and the model reduces to Eqs. (1)-(3).

**Figure S1:**
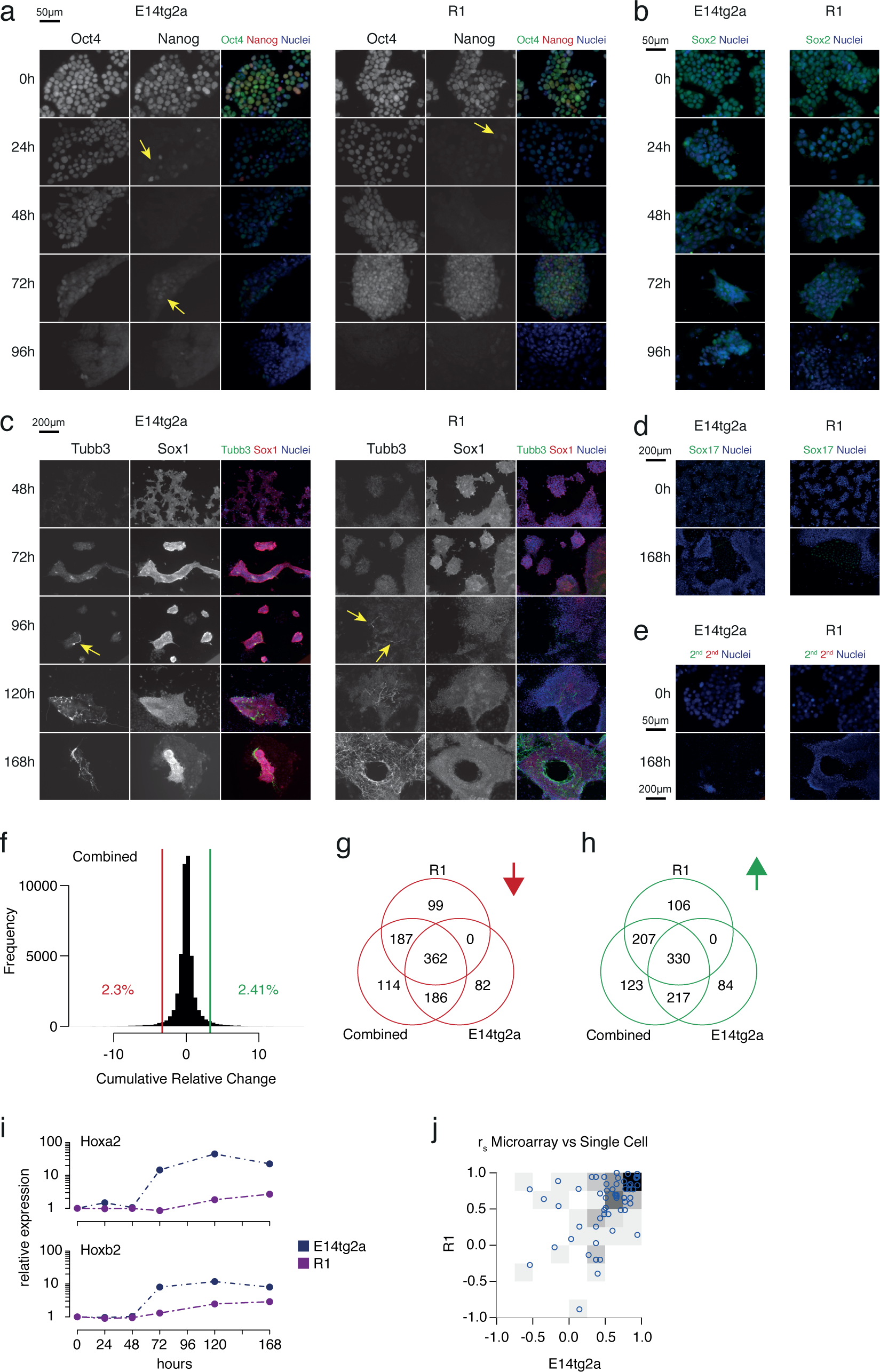
(**a-e**) Protein expression dynamics of (**a**) pluripotency associated transcription factors Oct4 and Nanog and (**b**) Sox2, (**c**) neuroectoderm markers Tubb3 and Sox1, (**d**) endodermal marker Sox17. Arrows show residual cells expressing Nanog and pioneer neurons respectively (**e**) Negative control. (**f**) Histogram of microarray-based expression changes across both E14 and R1 cells. (**g,h**) Venn diagram of (**g**) down- and (**h**) up-regulated genes. (**i**) Relative expression changes of Hoxa2 and Hoxb2. (**j**) Rank-correlation coefficient of average single-cell versus ensemble cell expression for R1 and E14 cells over time.

**Figure S2:**
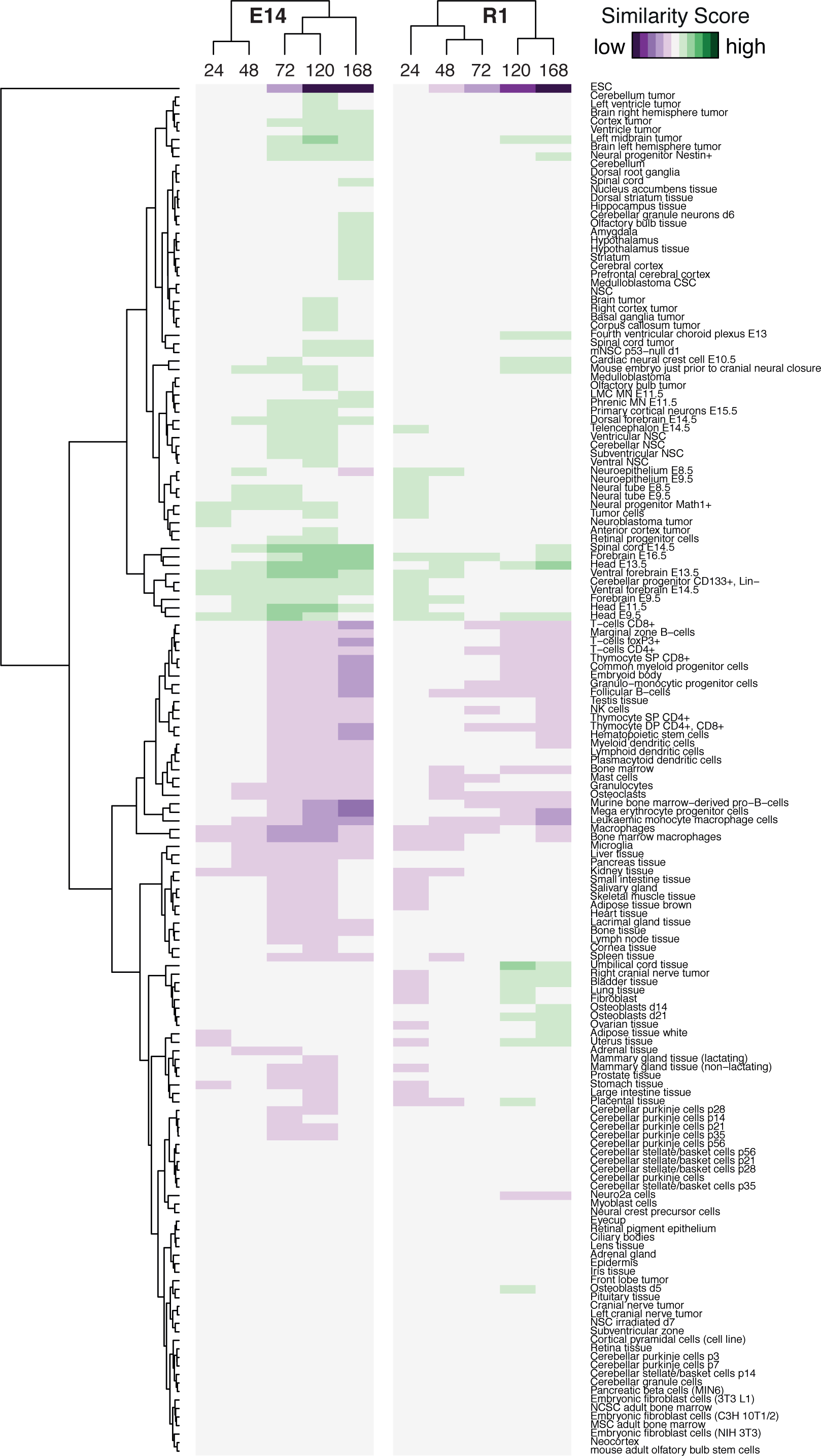
Similarity of time-course data for all of the 161 lineages we considered.

**Figure S3:**
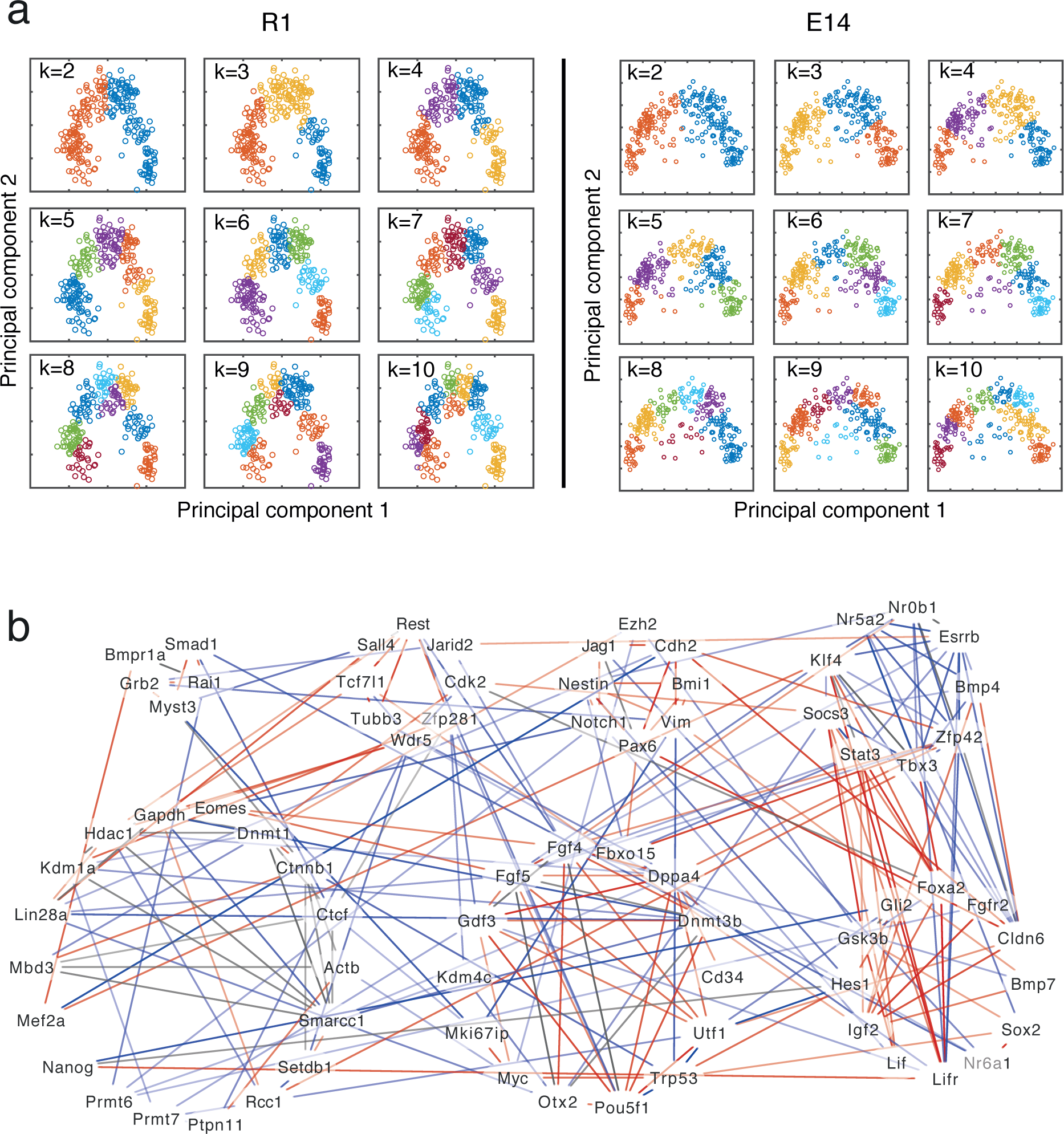
(**a**) PCA plot of k-means clustering with *k* ∈ {2,…, 10}. (**b**) (Co-)regulatory network inferred from single-cell data. Genes naturally grouped into seven modules based on an unbiased community detection algorithm (See main text for details). Significant interactions in cells classified as ESC or EPI are in blue, while significant interactions in cells classified as EPI or NPC are in red.

